# ChromaX: a fast and scalable breeding program simulator

**DOI:** 10.1101/2023.05.29.542709

**Authors:** Omar G. Younis, Matteo Turchetta, Daniel Ariza Suarez, Steven Yates, Bruno Studer, Ioannis N. Athanasiadis, Andreas Krause, Joachim M. Buhmann, Luca Corinzia

## Abstract

**Summary:** ChromaX is a Python library that enables the simulation of genetic recombination, genomic estimated breeding value calculations, and selection processes. By utilizing GPU processing, it can perform these simulations up to two orders of magnitude faster than existing tools with standard hardware. This offers breeders and scientists new opportunities to simulate genetic gain and optimize breeding schemes.

**Availability and Implementation:** The documentation is available at https://chromax.readthedocs.io

The code is available at https://github.com/kora-labs/chromax

## Introduction

Livestock and plant breeding is crucial to sustainable agriculture (Schön and Simianer, 2015) and to develop new breeds more suited for specific environments or market demands (Qaim, 2020). Recently, the availability of genomic data and advanced statistical methods have revolutionized breeding programs (Kim et al., 2020). Notably, genomic selection allows a breeder to predict the performance of an individual based on genetic makeup, avoiding expensive phenotyping (Meuwissen et al., 2001; Crossa et al., 2017). These new methods unlock various design possibilities for breeding schemes, making it harder to optimise them. Moreover, a single breeding cycle can take many years, involving many design choices during the process. Thus, there is a growing interest in using simulations to optimize breeding programs. The existing tools for simulating crosses are implemented in R (Broman et al., 2003; Mohammadi et al., 2015; Gaynor et al., 2020; Pook et al., 2020) or Julia (Chen et al., 2022). While they provide an extensive set of features, they are not capable of exploiting parallelism in high-performance computers that may be necessary for large and complex breeding schemes. For example, simulating a full-diallel cross of ten individuals with ten offspring results in 450 offspring, while a similar diallel of 20 individuals generates 1,900 offspring. With this rapid scaling, simulating a full diallel in a breeding program with thousands of individuals may be unfeasible; and hence developing tools that can speed up simulations is desirable. The most attractive language for this purpose is Python. Python is one of the most used programming languages for numerical computing and data science, with many libraries available for optimization and machine learning (Harris et al., 2020; Virtanen et al., 2020; Bradbury et al., 2018; Pedregosa et al., 2011; Paszke et al., 2019). Some of them allow exploiting parallelization capabilities of specialized hardware devices, such as GPUs and TPUs. Thus, developing a parallelizable Python tool for simulating crosses can increase the throughput of simulations and opens new opportunities to improve breeding efficiency. In this paper, we introduce ChromaX, a fast and scalable Python library that enables the stochastic simulation of the most common features in a breeding program like genetic recombination, fixation of genomes by doubled haploid induction, and selection. It can further calculate the Genomic Estimated Breeding Value (GEBV), and simulate genotype-by-environment interactions. ChromaX has been designed and implemented with scalability in mind, and exploits recent advances in that direction from the Python language.

### Software description

ChromaX is based on the high-performance numerical computing library JAX (Bradbury et al., 2018). Using JAX, ChromaX functions are compiled in XLA (Accelerated Linear Algebra) (Sabne, 2020), a compiler for linear algebra that accelerates function execution according to the domain and hardware available. This allows ChromaX to run seamlessly on various devices, such as CPUs, GPUs, and TPUs exploiting the parallelization offered by a variety of high-performance computing devices. ChromaX is available with the open-source license 3-Clause BSD. Source code is available on GitHub and is also distributed via the Python package installer “pip” as *chromax*. In the following sections, we describe the core functions available with ChromaX; for a complete list see the documentation.

#### Genotypic data

The simulator is initialized by providing a genetic linkage map supplied as a Pandas DataFrame (McKinney et al., 2010) or a path to a spreadsheet. In the genetic linkage map, each row represents a marker and columns contain the chromosome identifier, the position of the marker in centimorgans, and a column for each trait containing linear marker effects. Instead of marker positions, it can include a column indicating the probability of recombination occurring after the marker and before the next. The argument *trait names* can be used to specify a list of trait names; every element must match a column name in the genetic linkage map. A heritability value for each trait can also be specified.

#### Population data

ChromaX represents the genome of an individual as an array of shape (*m, d*), where *m* is the total number of markers and *d* is the ploidy. Thus, a population of *n* individuals is represented as an array of shape (*n, m, d*). Please note that the *d* axis of the array separates the homologous chromosomes, which is important for downstream recombination. As a result, the genetic data must be provided in a phased format. One way to achieve this is by using a tool like Beagle (Browning et al., 2021) for individuals that are heterozygous. The simulator allows the user to load population data from a file and save it at any point of the breeding program simulation.

#### Genetic recombination

ChromaX simulates the genetic recombinations that take place during meiosis to create new haplotypes. For simplicity, we assume diploid species and the Poisson model for crossover interference (McPeek and Speed, 1995). The genetic recombination function receives as input an array of genetic markers of the *k* parent pairs, i.e. an array of shape (*k*, 2, *m, d*). The function performs *k* crosses between pairs of parents and produces offspring with shape (*k, m, d*). The use of a single genetic recombination function for all biparental crosses allows the function to be parallelized across several dimensions, namely the number of crosses *k*, the number of parents (2), and the ploidy number. Generalizations to autopolyploid species and to other models for crossover interference are left for future developments.

#### Differentiable genetic recombination

In ChromaX, we further develop a novel genetic recombination function that generalizes the one that takes place in biparental crosses. This function computes the parents of a cross by taking the weighted average of a population. With a weight vector of dimension (*k, n*, 2), it performs *k* crosses on a population of size *n*. Using the JAX *grad* functionality, the user can obtain the analytical gradient with respect to parents’ weights. This provides a continuous relaxation of the genetic recombination that can be used to optimize the crossing by gradient-based methods (Polak, 2012).

#### Doubled haploid lines

ChromaX can simulate the fixation of genomes by doubled haploid induction. The user can specify the number of offspring per individual *dh*; ChromaX generates a line from each individual of the population. With an input population of shape (*n, m, d*), the generated population will be of shape (*n, dh, m, d*). Like the genetic recombination function, the parallelization occurs over the number of lines *n*, ploidy *d*, and the number of individuals per line *dh*.

#### Traits

ChromaX computes the GEBV for additive traits of a population using the marker effects available in the genetic linkage map or drawn from a standard probability distribution (e.g., normal distribution). The marker effects are represented as an array of shape (*t, m*), where *t* is the number of traits and *m* is the number of markers. ChromaX computes the genomic value by performing a tensor contraction of the marker effect with the input population array of markers of shape (*n, m, d*). The arrays are multiplied along the *m*-axis and then sum reduced along the *m* and *d* axes. The result is an array of shape (*n, t*) containing the GEBV of the population for each trait. This operation is well-suited for GPUs due to their ability to efficiently perform multi-dimensional multiplication and sum reduction.

ChromaX can further model phenotypes that have a genotype-by-environment interaction component, as described in Faux et al. (2016). In genotype-by-environment interaction, an environment is simulated as a random variable drawn from a normal distribution; this value multiplies a random additive trait that we fix at the beginning of the simulation with the variance determined by the heritability.

#### Selection

ChromaX enables the user to select the best individuals from a population based on a user-defined score. The score function accepts a population marker of shape (*n, m, d*) as input and returns an array of *n* scores. Examples of scoring functions included in the software are breeding values, (Lande and Thompson, 1990), optimal haploid value (Daetwyler et al., 2015), and phenotype.

### Performance

The performance of ChromaX is compared to AlphaSimR (Gaynor et al., 2020), a popular breeding program simulator. We perform this comparison on three hardware settings representing typical simulation conditions: an Apple M1 8-core, a computing cluster CPU (*AMD EPYC 7742 64-Core*), and a commonly available GPU (*Quadro RTX 6000*). At present, JAX uses only the CPU when it runs on the Apple M1. Table 1 shows the computation times when simulating 1k, 10k, and 100k biparental crosses from a population described by 1,000 markers across ten chromosomes. We do not benchmark AlphaSimR on GPU as it cannot run on this hardware. On the CPU, the computation time scales linearly with the number of crosses for both simulators and ChromaX is three to four times faster than AlphaSimR for the tested sizes. In contrast, on GPU the computation time scales sub-linearly (10× the computation time for 100× more crosses), and ChromaX can be hundreds of times faster than AlphaSimR.

**Table 1.** Simulation time in ms as a function of the number of crosses for AlphaSimR and ChromaX on different hardware settings (CPU: CPU computer cluster AMD EPYC 7742 64-Core; M1: Apple M1 8-Core; GPU: Quadro RTX 6000). Reported are the mean ± standard deviation over 100 simulations.

|  | Number of crosses |  |  |
| --- | --- | --- | --- |
|  | 1k | 10k | 100k |
| **AlphaSimR CPU** | 150 ± 5 | 1352 ± 9 | 14821 ± 149 |
| **AlphaSimR M1** | 87 ± 2 | 867 ± 5 | 8825 ± 36 |
| **ChromaX CPU** | 47 ± 1 | 473 ± 9 | 5149 ± 94 |
| **ChromaX M1** | 15 ± 0 | 147 ± 4 | 2086 ± 22 |
| **ChromaX GPU** | 3 ± 0 | 6 ± 1 | 29 ± 2 |

As a further comparison, we implement the breeding program described in Gaynor et al. (2020) with both simulators and compare the results. This breeding program, typical for an inbred species, assumes an initial diallel *F*_0_ characterized by 100 QTL per chromosome and 21 chromosomes. *F*_1_ is obtained by performing random biparental crosses from *F*_0_. Then, we obtain homozygous lines using the doubled haploid technique. To obtain the final cultivars, we simulate visual selection on head rows and preliminary, advanced, and elite yield trials, where individuals are evaluated with increasing accuracy for smaller population sizes. Fig. 1 shows the evolution of the population breeding values during the program. While some differences are expected due to the stochasticity of the process, the values are similar for both simulators. Table 2 reports the simulation times when the population sizes at the different generations are multiplied by various scaling factors. The results are similar to Table 1 with ChromaX achieving around 500*×* speed up on GPU compared to AlphaSimR.

**Table 2.** Simulation time in seconds for the inbred schema from Gaynor et al. (2020) as a function of the population size on different hardware. Mean values ± standard deviation over 100 simulations are reported.

|  | Scaling factor |  |  |
| --- | --- | --- | --- |
|  | 1x | 5x | 20x |
| **AlphaSimR CPU** | 3.37 ± 0.18 | 26.59 ± 0.34 | 95.71 ± 0.44 |
| **AlphaSimR M1** | 2.08 ± 0.02 | 16.64 ± 0.07 | 62.43 ± 0.21 |
| **ChromaX CPU** | 1.28 ± 0.02 | 6.36 ± 0.06 | 24.08 ± 0.31 |
| **ChromaX M1** | 0.34 ± 0.00 | 2.47 ± 0.05 | 14.04 ± 0.43 |
| **ChromaX GPU** | 0.02 ± 0.00 | 0.05 ± 0.00 | 0.15 ± 0.00 |

**Fig 1.**
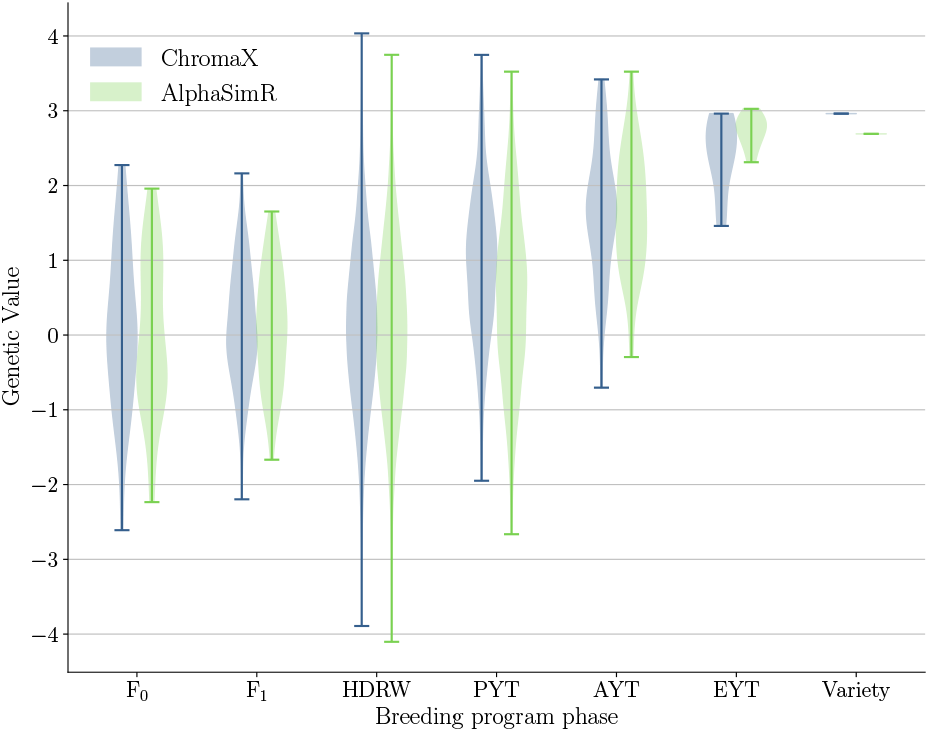
Comparision between the genetic value simulated by ChromaX and AlphaSimR for the same breeding schema. The F_0_ population contains 50 lines and F_1_ is created by 200 random biparental crosses. From each line in F_1_, 100 doubled haploids are obtained and evaluated in head rows (HDRW) using visual selection (low accuracy). Then plants are evaluated with increasing accuracy while reducing the population size. PYT: Preliminary Yield Trial; AYT: Advanced Yield Trial; EYT: Elite Yield Trial.

## Discussion

Our benchmarking experiments show that ChromaX is orders of magnitude faster in simulating sample breeding programs compared to existing software. Crucially, this will pave the way for the systematic exploration and optimization of complex designs in modern breeding programs.

## Limitations

ChromaX can simulate standard breeding programs but has limitations. First, ChromaX requires a genetic linkage map with marker effects and genetic data of the populations to simulate breeding cycles. To circumvent these requirements, other programs can simulate these data *in silico* by making some assumptions about the genetic features of the species. Second, ChromaX models additive traits and genotype-by-environment effects to simulate population phenotypes. ChromaX does not yet implement other biological effects, such as dominant or epistatic. Finally, ChromaX is designed for breeding programs of self- or open-pollinated species; other systems, such as hybrid breeding from distant heterotic groups, or monoecious/dioecious reproductive systems are not supported. Addressing these limitations will enable widespread use in a wider community.

## Acknowledgments

This work was financially supported by the Wageningen University and Research research theme “Data Driven Discoveries in a Changing Climate”.

